# Distinct anatomical and functional corticospinal inputs innervate different spinal neuron types

**DOI:** 10.1101/2024.04.29.591683

**Authors:** Samaher Fageiry, Claire L. Warriner, Jackson Loper, Liam Pianski, Thomas Reardon, Thomas M. Jessell, Rui M. Costa, Andrew Miri

**Author notes:** Correspondence (R.M.C.), (A.M.).

## Abstract

The corticospinal tract exerts its influence on movement through spinal neurons, which can be divided into types that exhibit distinct functions. However, it remains unknown whether these functional distinctions are reflected in the corticospinal inputs that different types of spinal neurons receive. Using rabies monosynaptic tracing from individual neuron types in the cervical cord and 3D histological reconstruction in mice, we discovered that different types receive inputs distinctly distributed across cortex, and aligned with cell type function. This included a distinct, sparse distribution of direct inputs from cortex onto motor neurons. Coupling rabies tracing with activity measurement during motor behavior revealed different interneuron types receive different input activity patterns, primarily due to the topographical distribution of the corticospinal neurons contacting them. Our results establish that different spinal neuron types get distinct anatomical and functional inputs from the cortex, and reveal functionally relevant homology to primate corticospinal organization.

## INTRODUCTION

Skilled movement is achieved by recruiting neural circuits distributed throughout the nervous system. Ultimately, these circuits impinge on spinal networks that engage motor neurons to drive task-appropriate sequences of muscle contraction. Although spinal circuits are capable of generating many basic motor patterns autonomously^1^, their activation in the context of more complex voluntary behaviors is dependent on inputs emanating from supraspinal networks in the cortex and brainstem^2,3^. It has long been appreciated that projections from deep layer cortical neurons to spinal cord via the corticospinal tract play a role in the control of skilled volitional movements such as grasping^4^ and visually guided locomotion^5^. These corticospinal projections originate broadly from multiple motor and sensory cortical regions and terminate throughout the spinal cord^6^.

One notable feature of corticospinal termination patterns is the conservation of direct contacts onto spinal interneurons across species. Layered onto this enduring characteristic is the emergence in some primates of direct contacts onto motor neurons; however, even in these primates the majority of corticospinal terminations are likely to be onto interneurons^7^. This suggests that conserved spinal interneuronal networks are critical recipients of descending corticospinal influence. This is reflected in the critical role spinal interneurons have now been shown to play in mediating corticospinal influence on grasping^8^ and in transforming descending motor commands into motor neuron activation patterns^9^. Despite this, many of the principles of organization and function of corticospinal projections onto interneurons have remained a mystery, in part because we had lacked the tools to delineate corticospinal neurons by their interneuron projection targets at cellular resolution.

Interneurons of the mature spinal cord are tasked with two principal roles, the relaying of sensory information from the periphery and the generation of motor output. The interneurons that subserve these functions are organized with a stereotyped topography along the dorsoventral axis, such that dorsal interneurons mediate sensory processing and ventral interneurons mediate motor processing^10^. Spinal interneurons can be further divided into discrete classes, each of which is marked by the expression of a distinct transcription factor^2,11^ and exhibits a distinct spatial distribution across the spinal gray matter^12^. The emergence of genetic strategies for interrogating the function of these types has led to the revelation of broad functional distinctions, despite the cooperative involvement of different types in the execution of particular behaviors, like locomotion^13–16^ and reaching^17,18^. Thus, spinal interneuron types exhibit some modularity in their functions, but we have yet to understand whether their functional distinctions are reflected in differences in the corticospinal inputs that they receive.

In this study, we characterized corticospinal inputs onto distinct spinal interneuron types, comparing their topography and temporal activity patterning. We focused on three different interneuron subclasses that reflect the rich functional repertoire of the spinal cord: (1) premotor excitatory interneurons that express the transcription factor Chx10, which derive from the V2 domain in the developing cord and include interneurons that mediate left-right coordination during locomotion^19^ and propriospinal function critical to reaching^20^; (2) post-sensory excitatory interneurons that express the peptide somatostatin (SST), which are critical players in the spinal circuitry underlying mechanosensation^21,22^; and (3) presynaptic inhibitory interneurons that express glutamic acid decarboxylase 2 (Gad2), which arise from the dI4 domain, are marked by expression of the transcription factor Ptf1a^23^, and modulate signaling at proprioceptive and cutaneous afferent terminals. The spinal location of their cell bodies differs according to their function: Chx10 cell bodies reside in the intermediate zone, Gad2 cell bodies occupy both the intermediate zone and dorsal horn, and SST cell bodies are primarily located in the superficial dorsal horn.

Using rabies transsynaptic tracing from individual types and 3D histological reconstruction, we have found that each of these spinal types does not sample proportionally from the overall spatial distribution of corticospinal neurons. Instead, they each receive distinctly distributed inputs aligned with their functions. Primary motor cortex was the only region to project abundantly to every interneuron type. Tracing from spinal motor neurons, we identified direct corticomotoneuronal inputs to forelimb-innervating motor neurons that arose from a much smaller cortical domain, almost entirely within primary motor cortex. Using an approach we developed to measure activity from corticospinals that contact specific interneuron types^24^, we establish that types can receive different input activity patterns resulting primarily from the regional distribution of corticospinals contacting them. We do find evidence for additional determinants of differing input to neuron types beyond this regional distribution, but the influence is comparatively small. Collectively, our results establish that broad-based but distinct corticospinal inputs drive interneuron types, and they expose functionally relevant homology to primate corticospinal organization.

## RESULTS

### Distribution of corticospinal projections

We first used retrograde viral tracing to comprehensively examine which cortical regions give rise to corticospinal neurons that engage forelimb spinal motor circuits. In adult mice (n = 4), AAV2-retro-GFP was injected into one hemicord throughout the dorsoventral extent of the spinal gray matter at spinal segments C4-T1 (Figure 1A). This led to robust labeling of corticospinal neurons in cortical layer 5B, primarily in the contralateral hemisphere (Figure 1B-D).

**Figure 1:**
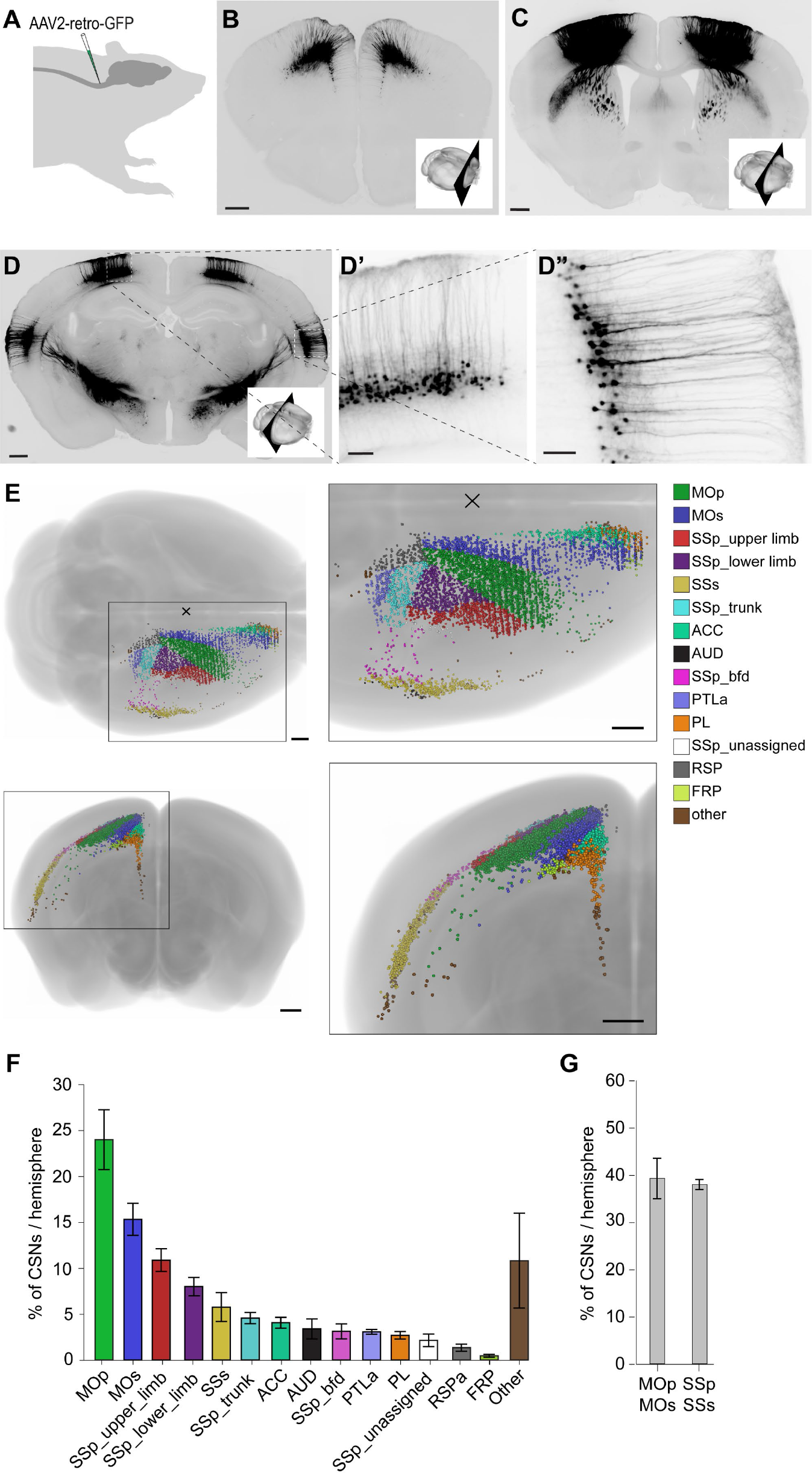
Distribution of corticospinal cell bodies innervating mouse cervical cord (A) Schematic illustrating the targeting of AAV2-retro-GFP to the cervical cord of adult wildtype mice. (B-D) Coronal histological sections showing backlabeled corticospinal neurons (scale bars=500µm; insets show rostrocaudal level of section). D’ and D” show magnified cell bodies at regions marked in D (scale bars=100µm). (E) 3D reconstruction of labeled corticospinal neurons in the contralateral hemisphere registered to the Allen adult mouse brain atlas. Cells are color coded by region. (F) Mean ± s.e.m. fraction of labeled corticospinal cell bodies localized to different cortical regions. (G) Mean ± s.e.m. fraction of labeled corticospinal cell bodies localized to primary and secondary motor cortex, and primary and secondary somatosensory cortex (right).

The cell body positions of corticospinal neurons in the contralateral cortex were mapped onto a 3D reference brain template^25^ (Figure 1E). We observed that primary (M1) and secondary (M2) motor cortex, contained 24.0 ± 3.25% and 15.4 ± 1.7% of the total labeled corticospinal cell bodies, respectively. Primary (S1) and secondary (S2) somatosensory cortex, contained 28.9 ± 2.5% and 5.8 ± 1.6%, respectively. Thus motor and somatosensory cortical regions each gave rise to similar proportions of corticospinal neurons (39% motor, 38% somatosensory), with S1 giving rise to more than any other individual region (Figure 1F,G). Corticospinal neurons observed outside of motor and somatosensory cortices were widely dispersed in multiple regions, including anterior cingulate (∼4% of all labeled corticospinal neurons), auditory (∼3%), posterior parietal association (3%), prelimbic (3%) and retrosplenial (∼1%) cortices.

Given the dorsoventral organization of spinal circuits by function, we next addressed which spinal circuits corticospinal neurons were likely to engage. We examined the termination locations of corticospinal neurons at cervical (C7), thoracic (T6), and lumbar (L3) levels to assess corticospinal engagement of upper-limb, trunk, and lower limb innervating circuits, respectively. *AAV2/1-Syn-hChR2-YFP* was injected in adult wildtype mice at several locations spanning forelimb M1 and S1 (caudal forelimb area^26^). Putative presynaptic corticospinal terminals were marked by their co-expression of YFP-tagged synaptophysin together with VGluT1 (Figure 2, top row). The position of these marked corticospinal terminals was normalized to reference coordinates in p56 adult mice and aggregated across animals (n = 3) in order to generate spatial density plots (Figure 2, bottom row).

**Figure 2:**
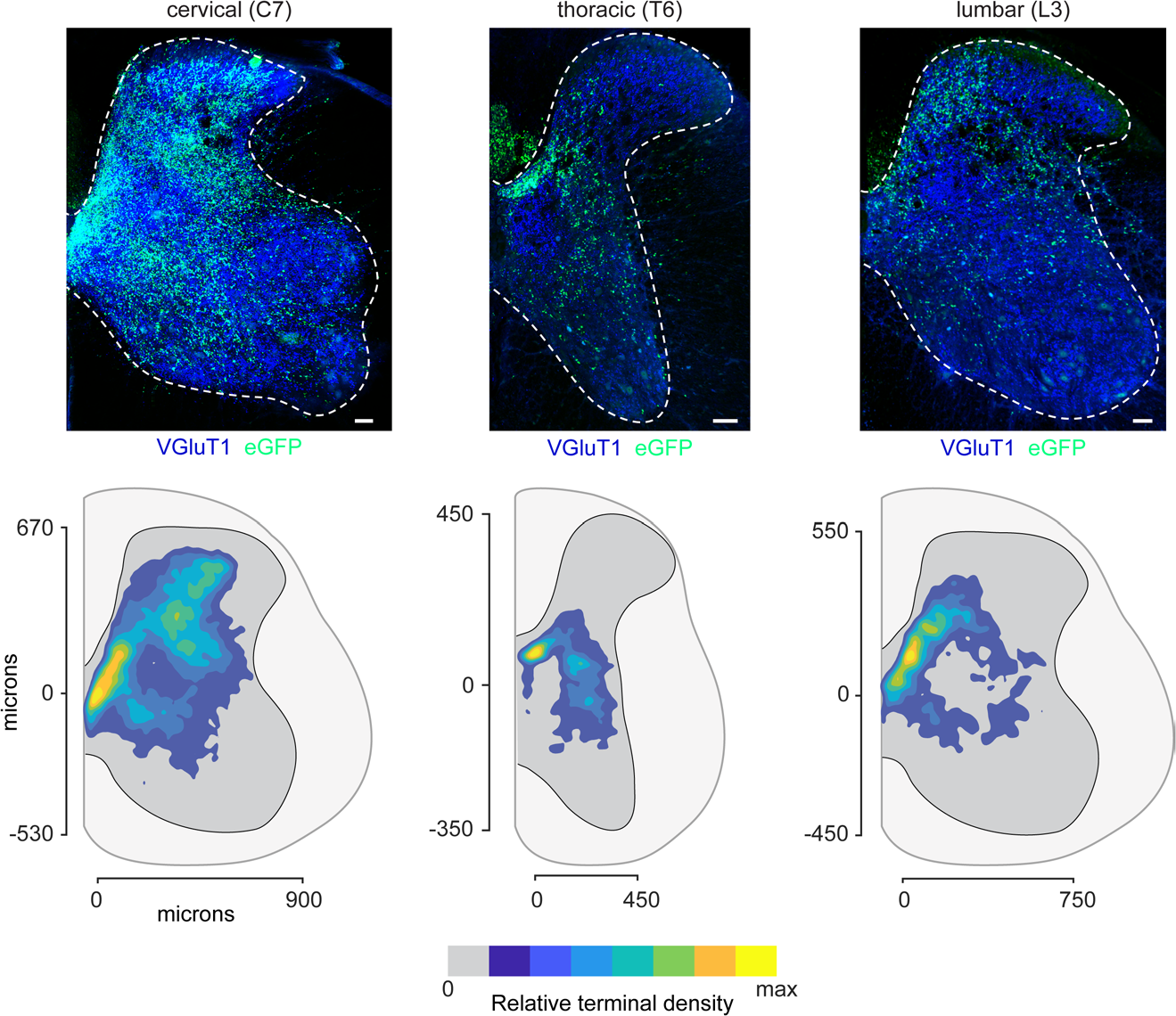
Corticospinal termination patterns at different rostrocaudal levels of the spinal cord (Top row) Fluorescence images of transverse spinal sections showing YFP labeled corticospinal terminals visualized in one hemicord after injection of *AAV2/1-Syn-hChR2-YFP* into contralateral motor and somatosensory cortex in adult mice (scale bars=100µm). Co-localization with VGluT1 represents putative presynaptic terminals. (Bottom row) Density maps of putative corticospinal terminations at each spinal level combined from three mice.

Corticospinal neurons exhibited the densest and broadest termination at cervical levels as compared to thoracic or lumbar. Cervical labeling was densest in the dorsal horn and intermediate zone but largely absent ventrally. A more restricted termination distribution was observed in the lumbar spinal cord where most labeling was observed on the medial aspect of the intermediate and deep dorsal horn. Corticospinal terminations were sparest at the non-limb innervating thoracic segments. In contrast, projections to the ipsilateral spinal cord were sparser and were concentrated in the inferomedial spinal gray matter and absent from the dorsal horn (Figure S1). Thus corticospinal neurons appear to engage both premotor and post-sensory spinal circuits, with the densest projections at cervical levels.

### Corticospinal connectivity with spinal interneuron types

The relative abundance of corticospinal terminations at cervical levels, coupled with the importance of neural circuits at this level in generating forelimb movement, led us to focus our analysis of postsynaptic corticospinal targets to these levels. We mapped the direct corticospinal inputs to Chx10^+^, Gad2^+^, and SST^+^ interneurons at these levels using rabies transsynaptic tracing. These interneuron populations were selected for their contrasting signatures with respect to somatic location, neurotransmitter production, connectivity, and function. Chx10 marks a population of glutamatergic premotor interneurons, SST marks a population of post-sensory interneurons and GAD2 marks a heterogeneous population of GABAergic presynaptic and postsynaptic inhibitory interneurons involved in sensory processing.

In order to visualize the monosynaptic corticospinal inputs to defined interneuron classes, we injected two Cre-dependent AAVs encoding the rabies glycoprotein, G, and the avian viral receptor, TVA (avian tumor virus receptor A), unilaterally into the cervical spinal cord, in three mouse strains where the expression of Cre was restricted to cells expressing Chx10, SST, or Gad2. This was followed two weeks later by an injection of pseudotyped rabies virus encoding tdTomato into the cervical spinal cord, thereby restricting the monosynaptic jump of rabies virus from those interneurons previously primed with G and TVA expression (Figure S2). The absence of G expression in presynaptic cells not otherwise in the starter cell population prevents the production of competent virus, and thus any further transsynaptic spread from these cells. At the injection site, those cells that were co-labeled with GFP and tdTomato represented starter cells from which the rabies virus jumped into monosynaptially coupled presynaptic neurons. The presynaptic population is labeled with tdTomato only. The position of individual presynaptic corticospinal neurons was visualized after histological sectioning and mapped onto a 3D reference mouse brain, as described above (Figure 3A,B; Figure S3).

**Figure 3:**
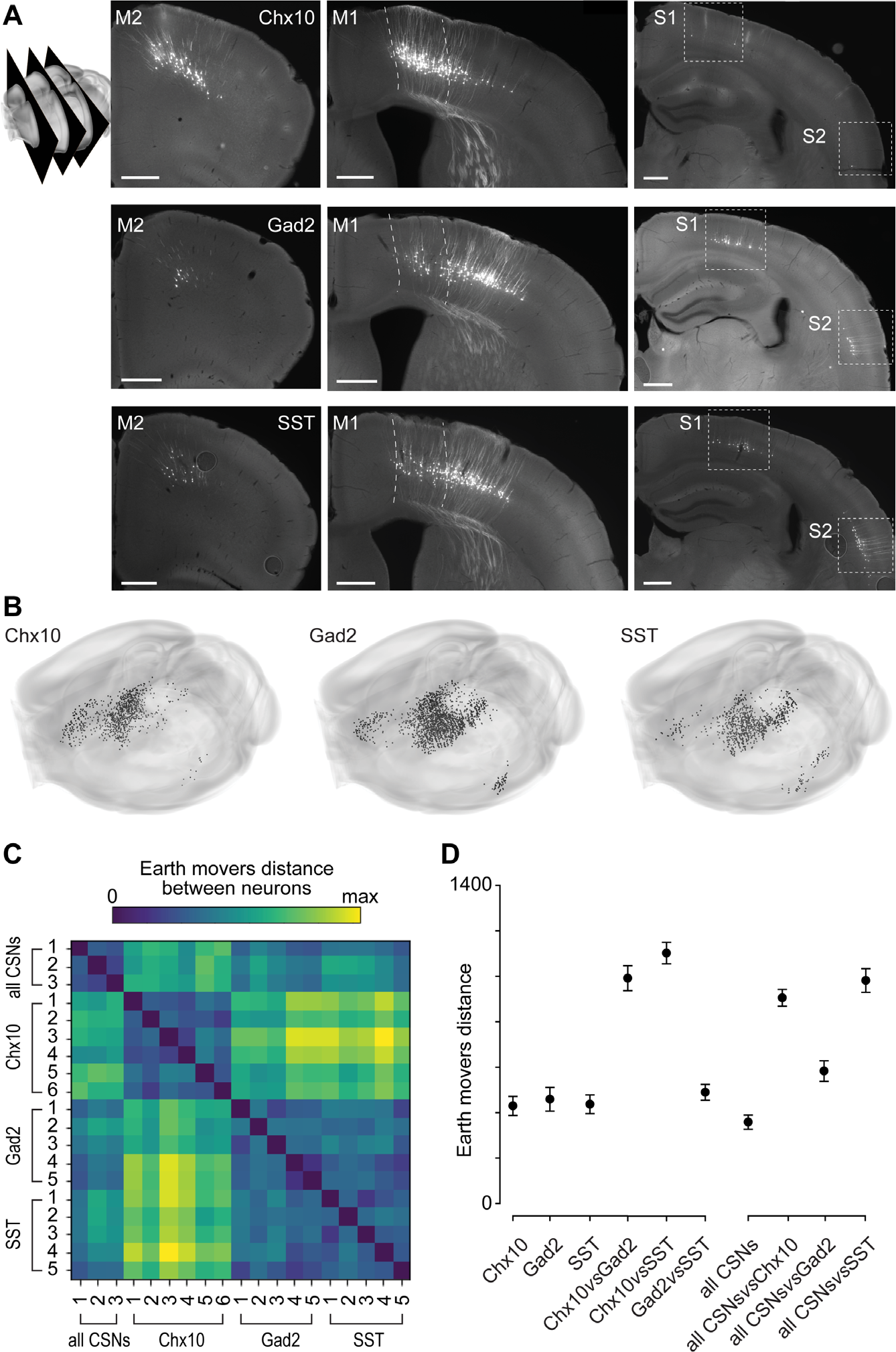
Variation in the spatial distribution of corticospinal cell bodies based on interneuron target (A) Example sections showing rabies tdTomato-labeled corticospinal neurons in the contralateral cortex following injection of pseudotyped rabies into cervical cord in Chx10::Cre, Gad2::Cre and SST::Cre mice (scale bars=500µm). Schematic at left depicts the rostrocaudal level of sections. (B) 3D reconstructions of the distribution of corticospinal cells in a single example mouse brain. (C) Earth mover’s distance (EMD) between spatial distributions of corticospinal cell bodies labeled from cervical cord in individual mice. Comparison includes the distribution of all corticospinal cell bodies using AAV2-retro-GFP (all CSNs) as well as monosynaptic labeling from Chx10, Gad2, or SST interneurons. Each row/column represents the distribution for an individual mouse. Mean ± s.e.m. EMD between spatial distributions of corticospinal cell bodies labeled from cervical cord in individual mice. All possible pairs of mice of each given type are included. The first three columns on the left plot reflect EMDs for mice of the same type (Chx10::Cre, Gad2::Cre, or SST::Cre). The first column on the right plot reflects EMDs for mice injected with AAV2-retro- GFP (all CSNs).

We observed abundant corticospinal neuronal labeling from Chx10, SST, and Gad2 spinal interneurons with most of the labeling residing in the cortical hemisphere contralateral to the injection site, as expected given that most of the corticospinal tract decussates. We observed a difference in the spatial distribution of corticospinal inputs to distinct interneuron populations; however, variability in the distribution of presynaptic labeling can arise from animal-to-animal variability or variability attributable to the genotype under study. To discriminate between the two, we calculated an earth mover’s distance (EMD) which quantitatively describes the element movements required to transform one distribution pattern into another (see Methods). A value of zero denotes that two distributions are identical, with higher values indicating greater distribution disparity. The variability between different mice of the same genotype was comparatively low (Figure 3C,D) and there was no statistically significant difference in the amount of variability between the genotypes (EMD for Chx10 429.4± 42.0, Gad2 458.5±152.3, SST 436.7±41.4 mean±s.e.m, p=0.585, one way ANOVA). In contrast the EMD was much higher when comparing the corticospinal input distributions of Chx10 and GAD2 interneurons or Chx10 and SST interneurons (Chx10vsGad2 p<0.001, Chx10vs SST p<0.001, one way ANOVA). The EMD of the inputs to Gad2 and SST interneurons, both of which are dorsally settled, is low and not significantly different from the EMD observed when comparing mice of the same genotype (p=0.102, one way ANOVA). Therefore, much of the difference between corticospinal inputs to these interneuron subsets can be explained by interneuron location in the spinal cord.

We also compared the disparity between the distribution of all corticospinal neurons (labeled using AAV2retro) and the distribution of corticospinals innervating specific interneuron subsets. Corticospinals innervating each subset were significantly more distant from the total corticospinal distributions (for Chx10 p<0.01, Gad2 p=0.04, SST p=0.018, one way ANOVA). In particular, Chx10-innervating corticospinals were the most unlike the total corticospinal population in terms of their distribution (Figure 3D). This is likely because there are very few Chx10-innervating corticospinals in posterior cortical areas (Figure 3B). These results indicate that each population of corticospinal neurons has a spatial distribution distinct from that of the overall population of corticospinals.

### Topographic variation in corticospinal connectivity

Most of the rabies-labeled corticospinal neurons were in either primary motor cortex (M1), secondary motor cortex (M2), primary somatosensory cortex (S1) or secondary somatosensory cortex (S2). These four regions accounted for 95% of all cortical labeling arising from Chx10 cells, 91% from SST cells, and 89% from Gad2 cells. We next asked whether the labeling in these four cortical sites differed among the spinal interneuron types studied (Figure 4A,B; note that from here on, percentages refer to the percent of total corticospinal neurons labeled in the contralateral hemisphere).

**Figure 4:**
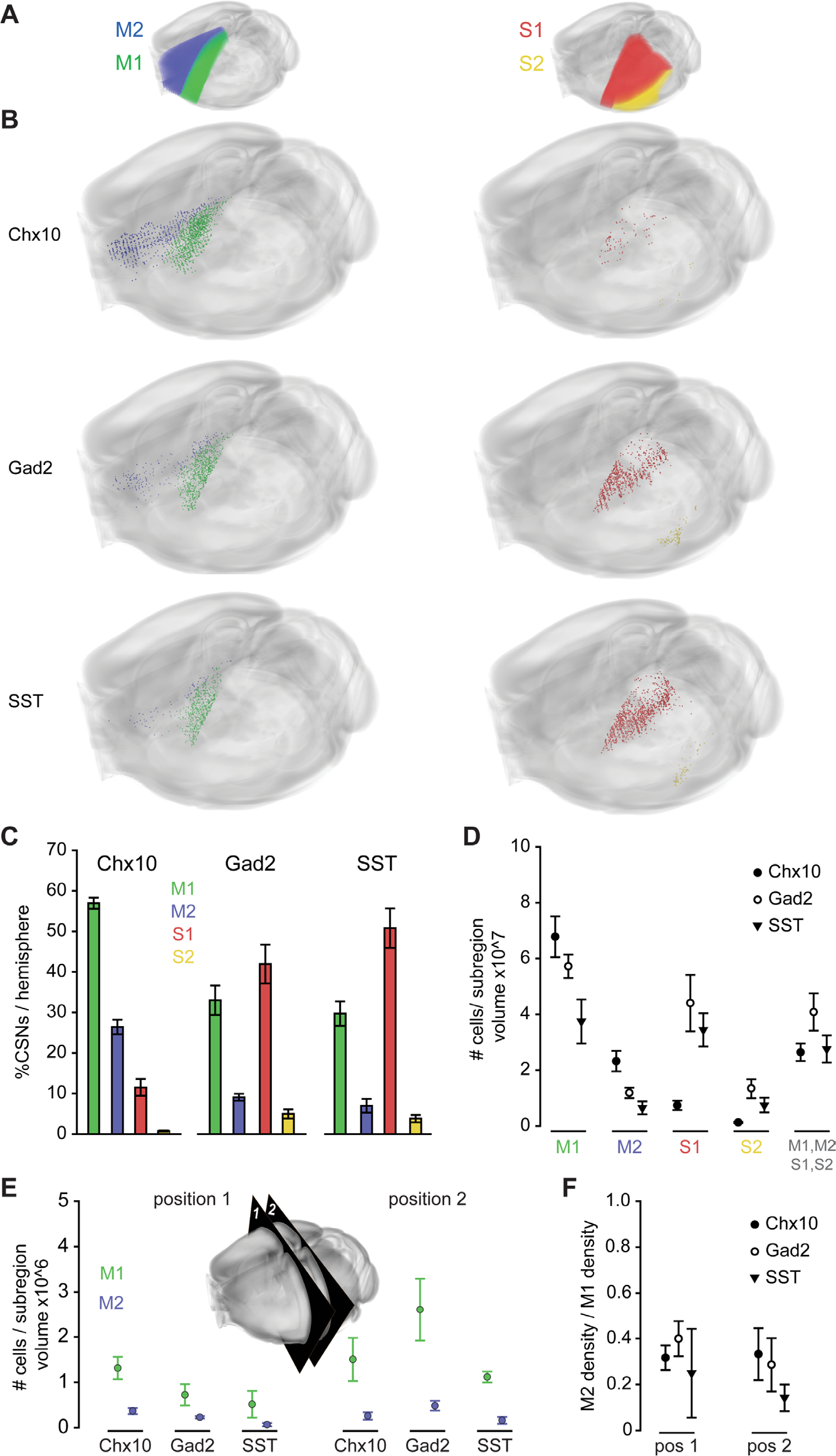
Variation in the regional distribution of corticospinal cell bodies based on interneuron target (A) Schematic depicting location of cortical regions as defined in the Allen Brain Atlas. (B) Examples of the distribution of corticospinal cell bodies labeled by monosynaptic tracing from Chx10, Gad2, and SST spinal interneurons in individual brains. (C) Mean ± s.e.m. percentage of corticospinal cell bodies located in M1, M2, S1, and S2 across animals. (D) Mean ± s.e.m. density of corticospinal cell bodies in M1, M2, S1 and S2 across animals, together with the total density over all four regions. (E) Mean ± s.e.m. cell body density in M1 and M2 across animals at both positions. Inset is a schematic depicting the two planes in which corticospinal cell body density in M1 and M2 were compared. Mean ± s.e.m. ratio of the cell body densities in M1 and M2 across animals at both positions shown in (E).

Each of the interneuron populations under study received substantial input from the primary motor cortex. M1 accounted for 56% of the cortically derived inputs to Chx10 interneurons, and 33% and 29% of corticospinal inputs to GAD2 and SST interneurons, respectively (Figure 4B,C Chx10vsGad2 p=0.034, Chx10vsSST p=0.008, Gad2vsSST p=1, one way ANOVA). In contrast, S1 accounted for the greatest proportion of corticospinal inputs to Gad2 and SST interneurons (41% and 50% respectively) but did not account for much of the input to Chx10 interneurons (11%; Chx10vsGad2 p<0.001, Chx10vsSST p<0.001, Gad2vsSST p=0.145, one way ANOVA). For all 3 populations, S2 accounted for ≤5% of the labeling and, as was the case for S1, SST and Gad2 showed more labeling than Chx10 cells (4%, 5% and <1% respectively Chx10vsGad2 p=0.009, Chx10vsSST p=0.049, Gad2vsSST p=1, one way ANOVA). Corticospinal neurons in M2 accounted for 26% of the retrograde labeling from Chx10 interneurons but were considerably sparser for GAD2 and SST interneurons, accounting for 9% and 7% of their labeling, respectively (Figure 4B,C ; Chx10vsGad2 p<0.001, Chx10vsSST p<0.001, Gad2vsSST p=0.375, one way ANOVA).

We next examined the density of corticospinal labeling in the four cortical regions across each interneuron type (Figure 4D). We observed variation in labeling density that for the most part aligned with interneuron type function. In M1 and M2, corticospinal neurons innervating Chx10 interneurons were the densest and those innervating more dorsal SST interneurons were sparser (in M1: Chx10vsSST p=0.021, Chx10vsGad2 p=0.287, Gad2vsSST p=0.132; M2: Chx10vsSST p=0.003, Chx10vsGad2 p=0.029, Gad2vsSST p=0.225, one way ANOVA). The opposite was observed in S1 and S2, where corticospinal neurons innervating SST and Gad2 were denser (S1: Chx10vsSST p=0.028, Chx10vsGad2 p=0.01, Gad2vsSST p=1. S2: Chx10vsSST p=0.05, Chx10vsGad2 p=0.005, Gad2vsSST p=1, one way ANOVA). Interestingly, however, we observed the corticospinals innervating dorsal interneuron populations were as dense in M1 as they were in S1, reflecting the overall prominence of input to each type from M1. These differences in density across the interneuron types innervated were more pronounced for individual regions than when combining all regions together (Figure 4D). Collectively, these results indicate differences across cortical regions in the distribution of inputs to the three interneuron populations examined, suggesting that cortical regions selectively engage interneuron types according to their function.

In examining the density of corticospinal neurons innervating each interneuron type, we found substantial differences in density across primary and secondary cortical regions (Figure 4D). In motor cortical regions, we found that corticospinals were consistently denser in M1 than in M2 for those innervating all three interneuron types. In somatosensory cortex, corticospinals were denser in S1 than in S2 for those innervating both dorsal interneuron types. For M1 and M2, we verified that this difference in density was not solely because of an overall difference in density along the rostrocaudal axis. Density differences were still observed within individual rostrocaudal sections (Figure 4E,F; position1: Chx10 p=0.015, Gad2 p=0.008, SST p=0.151; position 2: Chx10 p=0.047, Gad2 p=0.016, SST p<0.001, paired t-test or Mann-Whitney rank sum test). These differences in corticospinal density across regions could reflect differences in function across primary and secondary cortical regions, evidence for which remains limited in rodents^27,28^. In particular, these density differences may also reflect the partial hierarchy of cortical organization, in which both primary and secondary regions project to subcortical circuits, but primary regions exert comparatively more influence through these descending projections^28,29^.

### Direct cortical input to spinal motor neurons in the adult mouse

We next examined if there are direct corticospinal inputs onto spinal motor neurons in mice. Here we used a similar monosynaptic labeling approach as above. To selectively prime motor neurons for rabies labeling, we injected Cre-dependent AAVs into multiple forelimb muscles in ChAT::Cre mice at postnatal day 0. At adulthood we injected rabies virus into the ipsilateral cervical hemicord, imaged cortical sections, and aligned them to the reference brain (Figure 5A,B). Corticospinal labeling was observed, although in far fewer numbers than that seen following monosynaptic labeling from interneurons. The vast majority of corticospinal inputs to forelimb innervating motor neurons originated from M1, which accounted for 80% of all labeling. Though the function of these direct synapses onto motor neurons in rodents has not been established, this spatial distribution is very divergent from that of interneuron-contacting corticospinals and yet it parallels primate organization, where direct motor neuron contact is made by corticospinals heavily concentrated in the most caudal extent of M1^30^.

**Figure 5:**
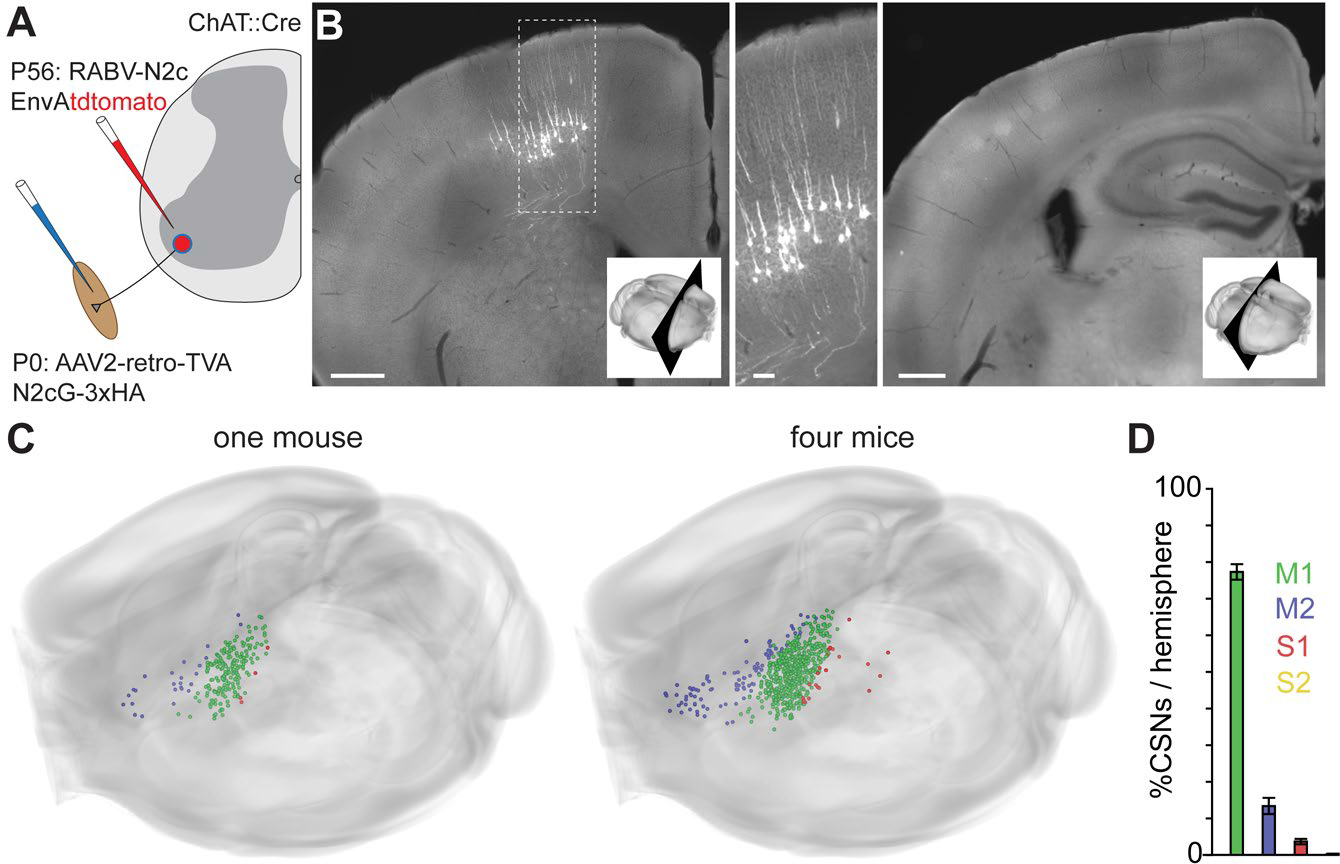
Rabies monosynaptic tracing from spinal motor neurons (A) At P0, AAV2-retro-TVA-N2cG-3xHA was injected into multiple forelimb muscles of ChAT::Cre mice to retrogradely prime forelimb innervating motor neurons for rabies monosynaptic tracing. In adulthood, RABV-N2CΔG-EnvA-tdTomato was injected into the cervical spinal cord (B) Fluorescence images of coronal brain sections showing rabies labeled corticospinal neurons in a ChAT::Cre mouse. Dotted outline shows region enlarged in middle image. Scale bars=500µm (left, right) or 100µm (middle). (C) Reconstructions showing the distribution of rabies labeled corticospinals in an example mouse (left) and all mice (right, n=4). Cells are color-coded by region according to the legend in (D). Mean ± s.e.m. percentage of motor neuron-innervating corticospinal neurons in different cortical regions.

### Distinct activity of cortical neurons that innervate different interneuron types

We next addressed whether corticospinal populations defined by their spinal interneuron type contact exhibit differing activity patterns during forelimb movement. To measure activity specifically in these corticospinal populations, we injected separate cohorts of Chx10::Cre and SST::Cre mice cervically with helper AAVs as above together with an rAAV2-retro that induces neuronal expression of GCaMP6f^31–33^ (Figure 6A). Mice were trained to perform a forelimb task under head-fixation in which they iteratively turn a small wheel with their right forelimb for water rewards^24^ (Figure 1B). The activity of four forelimb muscles (extensor and flexor pairs at both the elbow and wrist) were recorded using chronically implanted electromyographic (EMG) electrodes^34,35^. After mastering the task, mice were implanted with cranial windows over left frontodorsal cortex, and GCaMP6f fluorescence was measured during behavior with two-photon imaging. Fluorescence was measured in cross-sections of the apical trunk dendrites of labeled corticospinal neurons 350 mm below pia^36,37^ (Figure 6C,D). In each mouse, we performed several daily imaging sessions in various locations across S1, M1, and M2. We then injected rabies virus cervically, and green and red fluorescence image stacks were later collected from each imaging location to identify rabies labeled cells from which activity was measured^24^.

**Figure 6:**
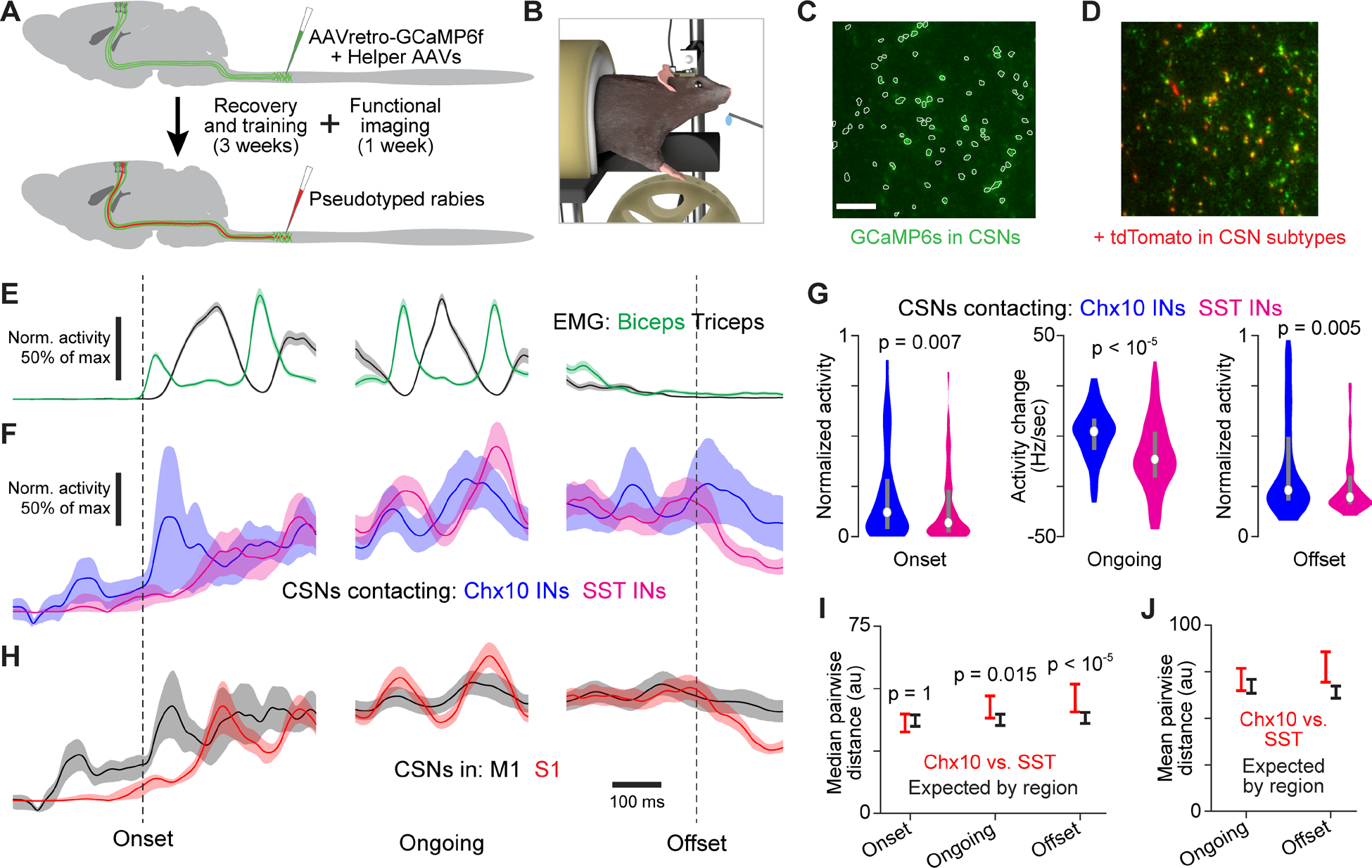
Corticospinal activity during single-forelimb wheel turning (A) Schematic of experimental timeline for measuring activity in corticospinal neurons, some of which could later be classified using rabies-driven tdTomato labeling. (B) Illustration of the head-fixed wheel-turning task. (C) Example of dendritic cross-sections from an imaging region 350 mm below pia. Scale bar=50µm. CSN, corticospinal neuron. (D) Overlay of GCaMP6f and tdTomato fluorescence for the window shown in (C). (E) Mean ± s.e.m. normalized muscle activity across trials for three behavioral epochs. (F,H) Mean inferred spiking across trials for corticospinal populations defined by spinal interneuron contact (F) or cortical region (H) during three behavioral epochs. Error bars reflect a 90% confidence interval from bootstrap resampling of imaged neurons. (G) Distributions across recorded neurons for the normalized, baseline-subtracted, trial-averaged inferred spike rate in the 50 ms following movement initiation (left) or the 200 ms following movement offset (right), and its rate of change in the 50 ms preceding peak triceps activity (center). P-values in G, I and J result from two-sided Wilcoxon rank sum tests. (I) 90% bootstrap confidence intervals for the median Euclidean distance between trial-averaged inferred spiking vectors for all possible pairs of neurons drawn one each from the two populations, compared to those expected based on regional location. 90% bootstrap confidence intervals for the mean Euclidean distance between trial-averaged inferred spiking vectors for all possible pairs of neurons drawn one each from the two populations, compared to those expected based on regional location.

To facilitate comparison of activity across corticospinal populations throughout movement execution, we examined activity during different epochs of the wheel-turning behavior. We estimated calcium fluctuations and spiking for identified cross-sections using established methods^38^ (Figure 6D). Using muscle activity recordings, we identified sets of similar behavioral segments (‘trials’) at the initiation of movement, during iterative turning, and at the offset of movement (Figure 6E). We observed differences in the trial-averaged activity of Chx10 and SST interneuron-contacting corticospinals in aggregate (Figure 6F,G). In particular, Chx10 interneuron-contacting corticospinals showed elevated activity at movement initiation, activity fell off faster at movement offset in SST interneuron-contacting corticospinals, and the average phase of peak activity relative to muscle activity differed between corticospinal populations. However, we observed similar differences when examining trial-averaged activity for all corticospinal neurons in M1 or S1 (Figure 6H). M1 corticospinals showed elevated activity at movement initiation, activity fell off faster in S1 corticospinals at movement offset, and the average phase of peak activity relative to muscle activity also differed.

We thus sought to assess the extent to which differences between the activity patterns of corticospinal populations could be explained by their cortical locations alone. To quantify the differences in activity patterns between neurons, we computed the Euclidean distance between trial-averaged activity for individual corticospinals during each behavioral epoch, treating each neuron’s trial-averaged activity from a given epoch as a vector. We computed confidence intervals for the median distance between trial-averages across all possible neuron pairs where neurons came from different interneuron-contacting populations (Figure 6I). We found that these distances were slightly but significantly higher than the equivalent distances for corticospinal neurons in M1 or S1. To quantify how much larger differences were for interneuron-contacting populations compared to the differences expected based on location, we computed the mean distance between trial-averages across all of the same neuron pairs. During ongoing wheel-turning, 85% of the distance between activity patterns is expected based on cortical location alone (Figure 6J). At movement offset, 93% of the distance between activity patterns is expected based on cortical location alone. This indicates that the activity patterns of corticospinals innervating different interneuron types are distinct, with the differences explainable mostly by the cortical region in which the neurons reside. This is consistent with the idea that the inputs to the corticospinal neurons in a specific location, rather than the neurons that they target, define their activity patterns.

## DISCUSSION

In this study, we investigated whether the corticospinal inputs to different spinal interneuron types are anatomically and functionally distinct. We found that corticospinal neurons that innervate the cervical cord have a broad distribution in cortex and are in general equally distributed between motor and somatosensory cortices. Furthermore, by selectively labeling synaptic inputs to three functionally distinct spinal interneuron populations, we discovered that the cortical populations that innervate them are largely distinct. Notably, we also detected sparser but clear direct contact onto spinal motor neurons from a more spatially restricted population of corticospinal neurons, the location of which resembles that seen in primates. Activity measurements during a single-forelimb motor task revealed that Chx10 and SST interneurons receive distinct functional signals from corticospinal neurons, though the differences can be explained largely by the balance of cortical regions the input comes from. Collectively, these findings indicate that spinal interneuron types receive broad-based but distinct corticospinal drive that is defined by the topography of the corticospinal neurons by which they are contacted.

### Cortical organization of spinal interneuron input

Our findings reveal that spinal interneuron types do not receive unbiased inputs from cortex proportional to the fraction of corticospinal neurons residing in each region. Instead, each interneuron type we examined selectively samples from the topographic distribution of corticospinals. Though we found that all three types receive abundant input from M1 corticospinals, M2 corticospinals preferentially innervate Chx10 interneurons and S1 corticospinals preferentially innervate SST and Gad2 interneurons. We note though that despite the selectivity of corticospinal sampling, this sampling is also broad-based, as each interneuron type received some degree of input from a broad range of cortical regions, spanning both sensory and motor. The similarity of the corticospinal distributions innervating SST and Gad2 interneurons, which have related but substantially different functions, also underscores the breadth of sampling. Cortical regions thus appear to broadly engage interneuron types, but with a proportionality that seems to mirror the degree of shared function. The graded nature of this topography between corticospinals and spinal interneurons is reminiscent of the recently described topography in cortical projections to medullary motor regions^39^.

Interestingly, we also observed a substantial difference in the balance of ipsilateral and contralateral contacts onto interneuron types. For Chx10 and Gad2 interneurons, 32% and 27% of labeled corticospinal neurons were ipsilateral respectively, while for SST interneurons, it was only 4%. It is known that corticospinal neurons originating from different cortical sites can differ in terms of the proportion of their connections that are ipsilaterally directed^40^. However, here we find a substantial difference in the ipsi-contra balance between SST and Gad2 populations that draw input from similar spatial distributions in the contralateral hemisphere. These balance differences we see may depend on the different spatial distributions of spinal interneuron types. Both Chx10 and Gad2 interneurons reside in the location of the spinal gray that has been shown to receive ipsilateral corticospinal terminations, whereas SST interneurons are chiefly distant to that site. We also note here that the sparseness of ipsilateral labeling from SST interneurons argues against bilateral spread of injected virus as a substantial cause of the apparent ipsilateral labeling from the Chx10 and Gad2 subpopulations.

We also found evidence that corticospinal input to spinal interneurons shows organization in terms of activity pattern. We observed that different interneuron types receive differently patterned input from corticospinals, and these differences go beyond what is expected based on differences in regional distribution of the corticospinals contacting each type. However, differences in regional distribution could account for the vast majority of the activity pattern differences. Thus for the types we examined here, the primary determinant of corticospinal input appears to be the regional distribution of cortical neurons from which they sample. Our activity measurements also exposed a substantial functional difference between M1 and S1 corticospinals. M1 corticospinals showed substantial activation just prior to and during movement initiation, while S1 corticospinal neurons were relatively quiet. The differences in the average phase of M1 and S1 activity during the alternating joint flexion and extension of wheel turning are also consistent with distinct motor and sensory functions. As we discuss below, much evidence suggests that corticospinal function is more biased towards sensory processing in rodents, relative to primates where involvement in motor control predominates. However, our activity measurements suggest that rodent M1 corticospinals targeting the cervical cord are primarily involved in driving movement. This contrasts with recent findings regarding corticospinal influence on the lumbar cord^41^.

### Relation between corticospinal topography and spinal interneuron function

Though we found that spinal interneuron types each receive broad-based corticospinal input, the balance of motor and somatosensory cortical inputs did align with the known functions of each type. We found that Chx10 interneurons receive direct corticospinal input, more than 80% of which originates from M1 and M2. Some Chx10 interneurons play a role in coordinating motor output across the right and left side of the body through their contact with commissural spinal interneurons. Another large subset of Chx10 interneurons also has a propriospinal character: these cells have a bifurcating axon, with one branch contacting local spinal motor neurons and the other contacting cells in the brainstem lateral reticular nucleus that relay propriospinal information to the cerebellum^42^. The blockade of propriospinal interneurons in the monkey impairs their ability to generate grasping movements in an attempt to retrieve food^8^. Transection of propriospinal axons in cats impairs reaching and food retrieval^43^. In mice, a global ablation of all Chx10 expressing neurons results in a reaching deficit^20^.

We found that corticospinal input to Gad2 interneurons arises from both motor and sensory cortical sites in almost equal proportion. A large subset of these interneurons mediate presynaptic inhibition at somatosensory afferent terminals, enabling them to dampen sensory feedback that may interfere with the execution of a motor task. Consistent with this capacity, selective ablation of cervical Gad2 interneurons in the mouse eliminated the usual smooth reach trajectory and unmasked a severe and reproducible forelimb oscillation during a skilled reaching behavior^18^. Motor but not somatosensory cortical areas have been shown to suppress spinal sensory inflow by activating presynaptic inhibition during the preparatory phase before movement^44^. However, subsequently during movement, somatosensory cortex did contribute to this engagement. The M1 and S1 corticospinal inputs to Gad2 interneurons we observed could mediate this engagement. The activity patterns of M1 and S1 corticospinals we observed are consistent with M1 involvement preceding movement and subsequent combined involvement of both populations.

We found that SST interneurons received their most abundant input from S1, just over 50% on average, consistent with top-down modulation of sensory input by S1 corticospinals. SST interneurons receive monosynaptic input from nociceptive primary sensory afferents, namely Aδ and C fibers, and contact ascending projection neurons in lamina I^45–47^. Ablation and optogenetic perturbation in mice have implicated SST interneurons in mechanical pain sensation and mechanosensory processing^21,22^. However, we found that SST interneurons receive a considerable proportion of their input from M1. This may suggest some shared function between motor and somatosensory corticospinals in the top-down modulation of sensory input.

### Comparative corticospinal organization

Though functionally relevant differences in organization exist, our findings revealed several general parallels between corticospinal organization in rodents, cats, and primates, suggestive of substantial homology across mammalian lineages. First, we found a significant contribution to the corticospinal pathway from somatosensory regions of mouse cortex (38%). This aligns with estimates of the fraction of the corticospinal tract that arises from parietal cortex in humans, monkeys, and cats (∼40%)^6^. Consistent with this, marked somatosensory deficits have been observed following lesioning of the corticospinal pathway in monkeys^48^ and mice^49^, reflecting a prominent and evolutionarily conserved role for the corticospinal tract in the modulation of spinal sensory circuits. Second, we found that M1 has a higher density of corticospinal neurons compared to M2, even after normalizing for rostrocaudal position. This is similar to the lower corticospinal density in premotor cortex compared to primary motor cortex in primates^50^ and cats^51^, and may underlie the higher current thresholds for evoking limb muscle activation from premotor areas^26,52^. Third, the ratio of M1 to S1 corticospinals contacting Chx10 interneurons is similar to the ratio of M1 to S1 corticospinals that contact forelimb motor neurons in monkeys, as previously observed with rabies-based transsynaptic tracing^30^. These observations suggest a cooperative involvement of M1 and S1 in limb motor control across species.

Lastly, using rabies labeling from cervical spinal motor neurons, we also observed direct cortico-motoneuronal synapses. Such connections have been observed in juvenile mice but appeared to be pruned away by adulthood^53^. Beyond development, such connections are generally thought to only exist in a subset of primates that rely more strongly on selective and independent digit use. Strikingly, we found that motor neuron-contacting corticospinals had a more restricted spatial distribution focused in caudal M1, paralleling observations in monkeys. These observations in monkeys have previously been used to suggest the emergence of a ‘new M1’ during primate evolution^30^. In interpreting our finding here, we stress several considerations. First, the number of motor neuron-contacting corticospinals we observed was much smaller than interneuron- contacting population sizes. Second, rabies could be traversing very weak or essentially dormant synapses, as previous electrophysiological studies in cats and rodents have found no evidence of appreciable cortico-motoneuronal connectivity^54^. Thus we do not interpret our observations as reflecting a substantial role for direct connections in mice, but rather only that “new M1” may have much earlier evolutionary origins.

Despite these similarities, our results reveal organizational distinctions between rodents and primates as well. In particular, we showed that SST interneurons receive a considerable proportion of their corticospinal input from M1 . SST cell bodies are principally located in the superficial dorsal horn, in correspondence with their sensory function. This contrasts with the observation in monkeys that M1 projections to the dorsal horn are generally sparse and completely absent from the more superficial laminae^55^. This SST connectivity, together with the prominence of dorsal and intermediate corticospinal terminations we found, align with the traditional view that the dominant involvement of corticospinal projections was originally in the control of sensory processing, with an increasingly motor-oriented role emerging across mammalian evolution^56^. Thus the abundant M1 connections to dorsal interneurons we found may reflect that the early and conserved role of the corticospinal pathway is to influence dorsal sensory circuits rather than its better studied role in motor control.

### Finer grained organization of corticospinal input to spinal interneurons

It is unlikely that individual corticospinal neurons are dedicated to contacting only a specific type of spinal interneurons. For example, we doubt that the similar spatial distributions of corticospinals that contact Gad2 and SST interneurons belie separate but spatially coextensive corticospinal populations. However, the methods employed here are not able to address this, as it would require demonstration of the efficacy of dual rabies monosynaptic tracing from two defined cell types into overlapping presynaptic populations in the same mouse. To the best of our knowledge, this has not yet been demonstrated. Emerging evidence suggests that motor cortical signals involved in driving muscle activity are broadly distributed across neurons^57^, suggesting that signals of distinct functionality may not be sequestered in specific neuronal subsets. This suggests that the bias we observe does not result from completely segregated connectivity between corticospinal populations and spinal interneuron types, but rather biases in such connectivity.

We did not examine if similar principles would hold for smaller subsets within the types we have analyzed here, which are each known to be functionally heterogeneous. Chx10 interneurons include both propriospinal interneurons as well as neurons whose projections stay within the spinal cord that help coordinate bilateral movements. Gad2 interneurons can be divided into at least two spatially restricted groups based on whether they project ventrally or dorsally^58^. Recent neurochemical and electrophysiological studies have revealed heterogeneity also among SST interneurons^59,60^. Moreover, recent evidence suggests a substantial degree of gene expression heterogeneity within spinal interneuron classes and subclasses^61–63^. It is possible that distinct subsets within a given class or subclass receive substantially different input patterns. We might expect though that this would similarly depend on selective sampling from the topographic distribution of corticospinal neurons.

## ACKNOWLEDGMENTS

We are grateful to E. Famojure, B. Han, K. Miao, Q. Liu, M. Mendelsohn, N. Zabello, K. MacArthur, M. Marshall, I. Shieren, G. Martins, H. Rodrigues, and M. Correia for technical assistance. Experiments were funded by an NIH BRAIN Initiative U19 (NINDS) 1U19NS104649 to R.M.C and T.M.J. S.F was supported by a Fulbright Science and Technology Scholarship. C.L.W. was supported by T32 and F31 NIH-NINDS training grants. A.M. was supported by a Searle Scholar Award, a Sloan Research Fellowship, a Simons Collaboration on the Global Brain Pilot Award, a Whitehall Research Grant Award, The Chicago Biomedical Consortium with support from the Searle Funds at The Chicago Community Trust, and NIH grant DP2 NS120847.

## AUTHOR CONTRIBUTIONS

S.F. and T.M.J. devised the project, designed experiments and secured funding with R.M.C. S.F., C.L.W., and A.M. performed the experiments and analyzed data. All authors helped interpret data. S.F. and A.M. wrote the manuscript, with input from R.M.C.

## STAR METHODS

### EXPERIMENTAL MODEL AND SUBJECT DETAILS

All experiments and procedures were performed according to NIH guidelines and approved by the Institutional Animal Care and Use Committee of Columbia University.

#### Mouse strains

Selective access to relevant neuronal populations for lineage tracing and virally mediated gene delivery was made possible by the use of the published mouse stains. A total of 110 adult mice were used, including those in early experimental stages to establish methodology (male and female for anatomical studies, males for behavioral studies). Strain details and number of animals in each group are as follows: Gad2::IRES::Cre mice (Jackson Laboratories stock #010802^64^); SST::IRES::Cre mice (Jackson Laboratories stock #018973); Chx10::Cre mice (L. Zagoraiou, S. Crone, K. Sharma, T.M. Jessell); Emx1::IRES::Cre mice (Jackson Laboratories stock #005628^65^); En1::Cre mice (Jackson Laboratories stock #007916^66^) and ROSA:: CAG::tdTomato mice (Jackson Laboratories stock #007914^67^) and C57BL/6J mice (Jackson Laboratories stock #000664). All Cre lines used were heterozygotes crossed to C57BL/6J mice (Jackson Laboratories stock #000664).

All mice used in experiments were individually or pair housed under a 12-h light/dark cycle. At the time of the measurement reported, animals were 8-1 weeks old. All animals were being used in scientific experiments for the first time. This includes no previous exposure to pharmacological substances or altered diets.

### METHOD DETAILS

#### Viruses

Table 1 lists the recombinant adeno-associated viruses (AAV) and rabies viruses (RABV) used in this study. The production of those viruses that were packaged within the Jessell lab is described below.

**Table 1.**
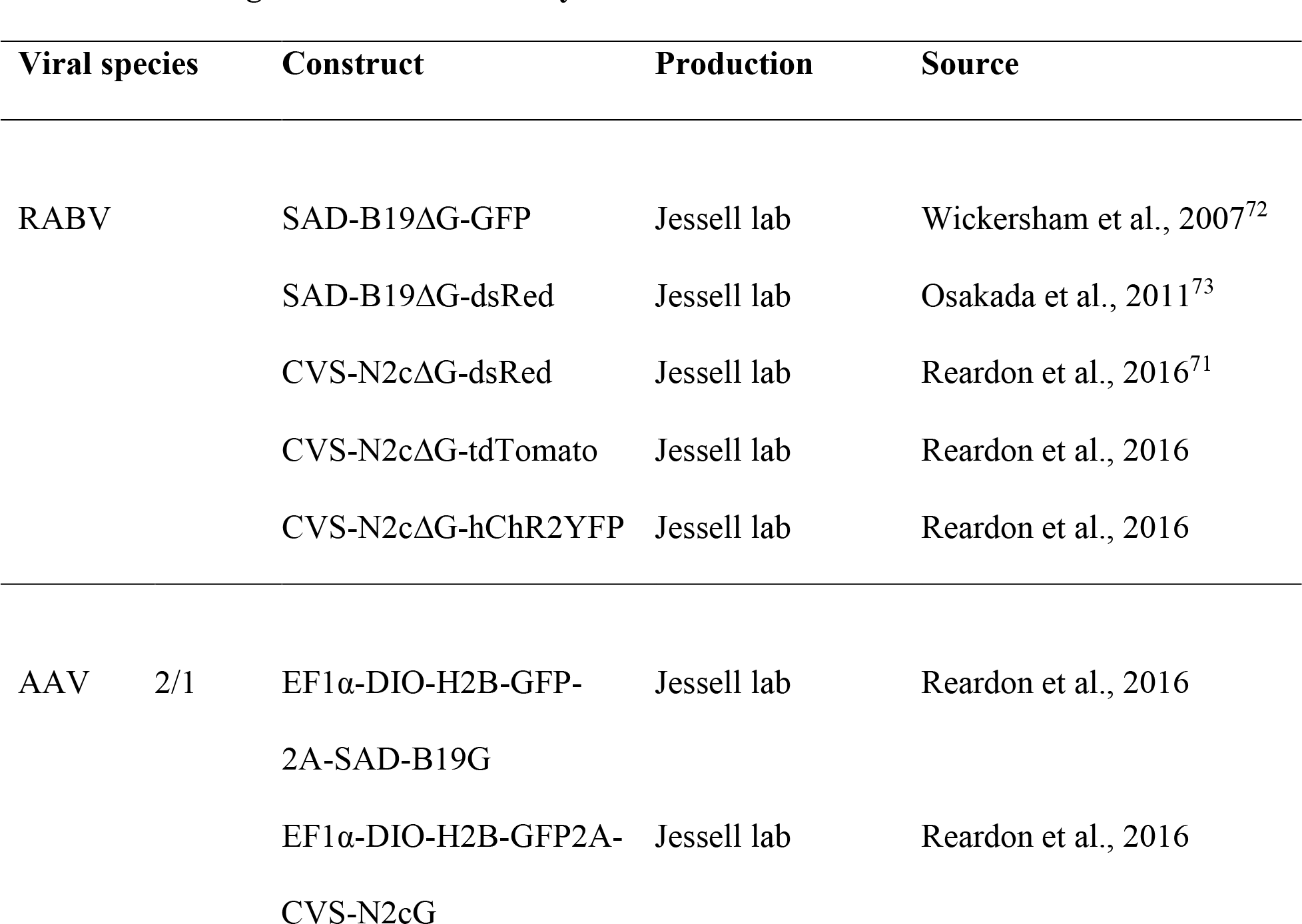
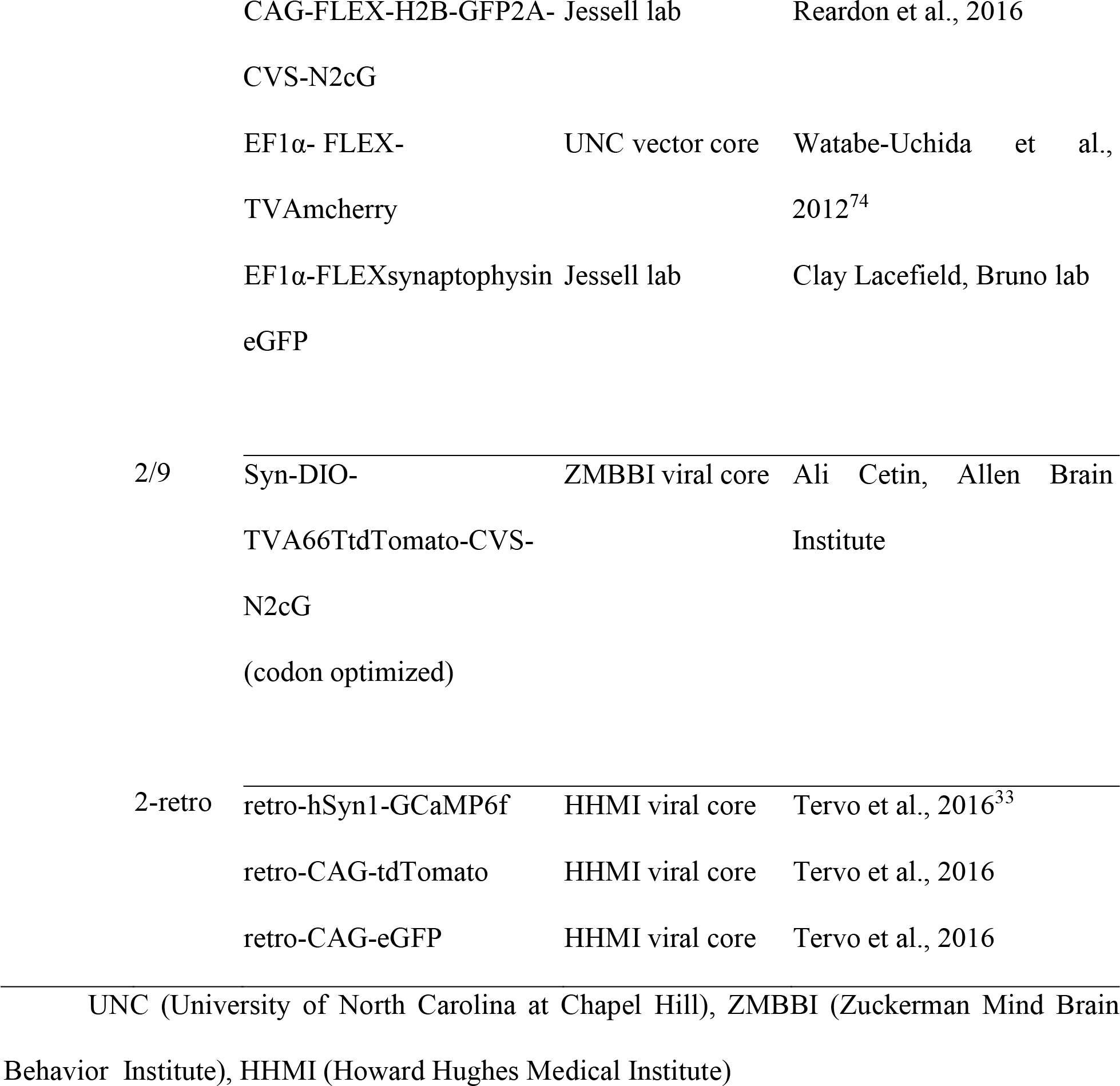
Viral reagents used in this study.

#### Production of chimeric AAVs

We used a previously described method of producing AAV vectors which combines the capsid proteins of AAV1 and AAV2 to generate a chimeric AAV that leverages the advantages of both^68^. Briefly, this involved calcium phosphate transfection of Hek293 cells with the AAV plasmid, the RepCap sequences of AAV1 (pH21) and AAV2 (pRV1) together with an adenovirus helper plasmid (pfΔ6). Cells were lysed 72 hours later to harvest the virions, which were subsequently purified by flowing the solution through a HiTrap heparin HP affinity column (GE life sciences). Finally, the AAV vector was concentrated using an Amicon ultra centrifugal filter with a 100,000 MW cutoff (Sigma-Aldrich).

#### Production of EnvA pseudotyped RABV

Amplification and pseudotyping of SAD-B19ΔG were performed as previously described^69,70^. Briefly, rescue stocks of wild type coat SAD-B19ΔG were amplified using BHK- B19G cells and the collected supernatant was added to BHKEnvA2 cells (cell lines courtesy of Ian Wickersham). The cells were gently washed with phosphate buffered saline (PBS) 24 hours later and again at 48 hours to remove unpseudotyped virions. Seventy-two hours after the second wash, the EnvA pseudotyped virus was collected, filtered using a 0.45 µm filter, and concentrated first by ultracentrifugation and then using an Amicon ultra centrifugal filter with a 100,000 MW cutoff.

The steps for producing EnvA pseudotyped CVS-N2cΔG closely mirrored those outlined above with a few notable exceptions. First, neuroblastoma derived Neuro2ACVS-N2cG and Neuro2A-EnvA cells were used in the amplification and pseudotyping steps, respectively. Second, cells were kept at 34°C and 3% CO2 to slow their metabolism and prolong survival. Finally, supernatant was collected over the course of at least one month during amplification and at least one week during pseudotyping^71^. Supernatants from the later process were pooled prior to ultracentrifugation.

#### Surgical procedures

Aseptic technique was applied throughout all surgical procedures. For all mice older than P7, anesthesia was achieved by Isoflurane (1-3%) and analgesia was delivered subcutaneously following induction and again prior to wound closure using carprofen (5 mg/kg) and bupivacaine (2 mg/kg), respectively. Hair overlying the surgical site was shaved and the mouse placed atop a warm heat pad maintained at 37°C. Eye ointment was then applied, and the incision site sterilized three times using betadine and ethanol (70%) soaked gauzes sequentially. Throughout the surgery the respiratory rate and effort of the mice was monitored together with the depth of anesthesia as assessed by the withdrawal reflex following hindlimb toe pinch. Closure of the incision was achieved using sterile sutures (nylon 6.0) and mice were subsequently observed continuously until they recovered from anesthesia. Postoperative analgesia was administered subcutaneously using carprofen (5 mg/kg) at 24 and 48 hours post-surgery and sutures were removed post-surgery unless the mouse was sacrificed before then.

#### Spinal injections

Cervical spinal surgeries were performed as previously described^17^ using a spinal stereotaxic frame (David Kopf Instruments). Briefly, reversal of the physiological lordosis of the cervical spine to expose the intervertebral spaces was achieved by first slightly tilting the head forward and subsequently elevating the tail using a spinal vertebral clamp. A vertical incision was made that extended from the occiput to a midline point extrapolated from the inferior tip of the scapulae. The exposed trapezius muscle was separated at the midline using forceps and the underlying musculature gently retracted on either side. Afterwards the muscles which insert into the vertebral column itself were gently removed using a delicate bone scraper (Fine Science Tools). To maximize separation of the cervical vertebrae and better expose the spinal cord, we clamped the very prominent T2 spinous process using a serrated clip attached to the stereotaxic frame. Finally, to expose the spinal segments, we removed the overlying dura using forceps or a microsurgical knife (Fine Science Tools).

The virus to be injected was loaded into a pulled glass capillary tube using a Nanoject II/III injector (Drummond Scientific) attached to the stereotaxic arm. The capillary tip was zeroed at the cord’s surface and gradually lowered to the ventral most injection site. Multiple injections of 13.8 or 23nl were made, retracting every 50 or 100μm, respectively. After the final injection, we waited one minute before fully retracting the capillary tip from the cord. For injections of AAV I injected at two sites in each brachial segment of the hemicord unilaterally from C4 to T1 in order to maximize coverage of the spinal gray matter. In experiments where EnvA pseudotyped RABV was used, the incision was reopened two weeks after AAV injection, and virus was injected into the same brachial segments (13.8nl boluses, three sites per segment).

#### Cranial injections

The skull was exposed by a midline incision and leveled so that bregma and lambda were in the same Z plane. We subsequently leveled in the horizontal direction so that corresponding points on the right and left side were in the same X plane. A burr hole overlying the target site was made with a dental drill, and a pulled glass capillary containing the desired injectant was lowered into position. Following the injection, we waited 5 minutes before slowly retracting the capillary. For cortical injections, we injected AAV2+1 EF1α-FLEX-synaptophysin eGFP at multiple sites into the rostral and caudal forelimb areas^26^.

#### Histology

Tissue from lineage tracing and viral tracing experiments was processed for histology following transcardial perfusion of deeply anesthetized mice. Following a thoracotomy and right atrial incision, cold phosphate buffered saline (PBS) followed by 4% paraformaldehyde (PFA)/0.1M PB was delivered into the left ventricle using a butterfly needle (Becton Dickinson). Brains were postfixed overnight in 4% PFA/0.1M PB at 4°, washed with cold PBS and subsequently mounted in 4% agarose for vibratome sectioning (100 µm). For vibratome sectioning of spinal cords (80 µm), we first carried out a ventral laminectomy to expose the underlying cord, post-fixed overnight, washed with PBS, removed the overlying meninges, and embedded in 4% agarose. For cry sectioning of spinal cords (19-40 µm) the post-fixation step was shortened to two hours. Following PBS washes, the cords were immersed in 30% sucrose/0.1M PB until fully equilibrated, then OCT (Tissue Tek) embedded and snap frozen on dry ice.

#### Immunohistochemistry

The following primary antibodies were used: rabbit anti-GFP (1:1000, Invitrogen), chicken anti-GFP (1:4000), rabbit anti-DsRed (1:100, Clontech), goat anti-ChAT (1:200, Millipore), rabbit anti-DsRed (1:100, Clontech) and guinea pig anti-vGluT1 (1:32000^23^). Secondary antibodies conjugated to Alexa-488, Cy3 or Cy5 were generated in chicken or donkey and diluted 1:500 (Jackson Immunoresearch Laboratories).

Cryosectioned or vibratome-sectioned spinal cords were incubated in primary antibody for one or two nights, respectively. Following PBS washes, cords were exposed to fluorophore-conjugated secondary antibodies for 3 hours at room temperature (cryosectioned tissue) or overnight at 4°C (vibratome-sectioned tissue). Images were acquired on a Nikon SMZ18 Stereo Imaging system or an LSM 710 Meta Confocal (Carl Zeiss).

#### Electromyographic electrode and headplate implantation

Electromyographic (EMG) electrodes were fabricated for forelimb muscle recording using previously described methods^75,76^. Briefly, electrode sets included four electrode pairs, one for each recorded muscle. Each pair consisted of two 0.001-inch braided steel wires (793200, A-M Systems) knotted together. The insulation was stripped from 1 mm to 1.5 mm away from the knot on one wire of each pair, and from 2 to 2.5 mm away from the knot on the other. The ends of the wires above the knot were soldered to a miniature connector (11P3828, Newark). The length of wire left between the knot and the connector was 3.5 cm for elbow muscles (biceps and triceps, lateral head, were implanted), and 4.5 mm for wrist muscles (extensor carpi radialis and palmaris longus were implanted). The remaining were twisted together and crimped within a 27-gauge needle to facilitate muscle insertion.

Surgical implantation procedures for electrode and headplate implantation have also been previously described^24^. Briefly, under anesthesia and following hair removal, incisions were made above the muscles to be implanted. Electrode pairs were guided from an incision at the scalp to forelimb incisions. Electrodes were inserted through targeted muscles using the needles. Wires were knotted just distal to muscle exit, excess wire and needle were cut away. During the same surgery, 3D-printed headplates^77^ were attached to the skull using dental cement (Metabond, Parkell). The EMG electrode connector was cemented to the posterior edge of the headplate. After recovery from the implantation surgery, mice were placed on a water schedule in which they received 1 mL of water per day.

#### Wheel-turning task

Male mice were trained through a behavioral shaping procedure to use their right forelimb to repeatedly pull the rungs of a small 3D-printed wheel. The experimental apparatus has been previously described^24^. Briefly, head-fixed mice sat within a curved hutch (4.5 cm outer diameter, 4.8 cm length) lined with soft foam (3 cm inner diameter) that covered the posterior half of their body. Their right forelimb rested on a wheel (60 mm diameter, 11.5 mm width) bearing 28 rungs (1.4 mm diameter), while their remaining limbs rested on a platform. A barrier below the mouse’s chest prevented access to the wheel by the left forelimb. A ratchet ensured that the wheel could only rotate in one direction. Experimental control was performed using the MATLAB Data Acquisition Toolbox and the NI PCIe-6323 DAQ. The wheel was mounted on a stainless-steel shaft, on which was mounted an absolute rotary encoder (A2K-A-125-H-M, U.S. Digital). Water rewards (3 μL/droplet) were dispensed by a solenoid valve (161T012, NResearch) attached to a feeding needle (01-290-12, Fisher).

Water-scheduled mice were acclimated to being handled by the experimenter and head-fixation using established protocols^78^. After two daily sessions of acclimation to handling, mice were acclimated to being head-fixed on the behavioral apparatus for 15 min on the first day and 30 min on the second day, during which time they were provided with water rewards (3 μL per reward) at regular intervals. During acclimation, the wheel was locked in place to prevent its movement.

After acclimation, mice underwent daily 45-min training sessions to master the wheel-turning task. During initial training sessions (1-2 sessions), the wheel was unlocked and the experimenter initiated rewards whenever the mouse touched the wheel or rotated it, even slightly. Mice gradually learned to associate wheel rotation with reward and began iteratively pulling the rungs of the wheel. Over the subsequent sessions (4-6 sessions), mice were trained via an automated training script to pull the wheel increasingly fast and increasingly far. During each bout of wheel-turning, the script integrated the wheel rotation distance. When a certain threshold was reached, the time to reach the threshold distance was recorded. Bouts during which the threshold was not reached were ignored. Rewards (3 μL water) were dispensed for each of the first 10 instances in which the distance threshold was met. Subsequently, the time to threshold was compared to that of the 10 most recent instances and rewards were provided according to the following scheme: a time to threshold below the 10th percentile value yielded four rewards; a time below the 40th percentile value yielded two rewards, and a time below the 75th percentile value yielded one reward. If the time was above the 75th percentile value, no rewards were dispensed. The rate of reward dispensation was computed every minute and adjusted to maintain the reward rate such that the mouse would receive ∼1 ml of water per session.

#### Two-photon imaging during behavior and EMG recording

Procedures for two-photon imaging followed those we recently published^24^. For these experiments, mice underwent cervical spinal injection surgery the day prior to EMG electrode and headplate implantation surgery. AAV2-retro-hSyn1-GCaMp6f (HHMI Viral Core) was injected along with helper viruses to drive neuronal expression of GCaMP6f.

After mice reached proficiency with the wheel-turning task, cranial windows were implanted on a rectangular craniotomy (∼1.5 mm mediolateral by 3 mm rostrocaudal) centered on the CFA following dura removal. Windows were constructed from two custom-shaped, laser-cut, truncated glass pieces (200 μm thick, Tower Optical Corp.) adhered to a larger 4 mm diameter (No. 1, Warner Instruments) glass coverslip using UV-curing adhesive (Norland Optical Adhesive 61). The cranial window was lowered into place held by suction through a glass capillary tube. The edge of the coverslip rested on the skull and was affixed using dental cement.

Two-photon fluorescence imaging was carried out using a 25×1.0NA objective (Olympus) mounted on an Ultima microscope (Prairie Technologies) and a Ti:sapphire laser (Chameleon Ultra II, Coherent) tuned to 940 nm (GCaMP6f) or 1050 nm (tdTomato). GCaMP6f image time series were acquired during wheel-turning behavior for 20 to 40 minutes epochs for up to 7 daily behavioral sessions. Each session targeted a different region of cortex. GCaMP6f image time series were acquired at a framerate of 60 Hz, and every four images were averaged, yielding 15 Hz time series. Imaging windows were ∼300 × 300 μm with 256 × 256 square pixels. EMG recordings were amplified and bandpass filtered (250–20,000 Hz) with a differential amplifier (MA102 with MA103S preamplifiers, University of Cologne electronics lab) and acquired at 10 kHz (General Purpose Input Output Box, Prairie Technologies).

After functional imaging was complete, RABV-N2CΔG-EnvA-tdTomato was injected. Ten days after rabies injection, Z image stacks were collected over 40–80 μm depths with 1 μm separating each image both for GCaMP6f and tdTomato in each plane. The red emission collection channel registered no detectable signal at 940 nm excitation, indicating that red emission was generated solely by tdTomato and not GcaMP6f.

### QUANTIFICATION AND STATISTICAL ANALYSIS

Statistical analyses of anatomical tracing experiments were performed in SigmaPlot. Error bars indicate the mean ± the standard error of the mean.

All analysis of functional imaging data was completed in MATLAB v.9.4–9.9 (MathWorks). Procedures for analysis of image time series followed those we recently published^24^. Details lacking in the summary below can be found in that paper. Briefly, EMG measurements for each muscle were high-pass filtered at 40 Hz, rectified, convolved with a Gaussian having a 10 ms standard deviation, and normalized to their respective 99.5th percentile value. GCaMP6f fluorescence image time series were motion-corrected with the ‘normcorre’ algorithm^79^. Individual dendritic cross-sections (sources) were defined from motion-corrected image time series using the CNMF function library^38^. Slow shifts in signal baseline or the resulting sources were corrected by identifying the 10th percentile value from each successive 25-sample time series segment and subtracting this value from samples within the given segment. Calcium fluctuations were estimated, and spiking was inferred using the ‘cont_ca_sampler’ and ‘make_mean_sample’ functions.

EMG recordings were used to identify segments to serve as trial equivalents. To find sets of similar segments during on-going wheel turning, peaks were found in biceps and triceps activity (Matlab function ‘findpeaks’). The modal durations from triceps to the previous and subsequent biceps peaks were found, using the peaks of histograms with 2.5 ms bins and ignoring durations < 50 ms. We then identified the set of at least 30 segments that were 400 ms in length and centered on a triceps peak, such that the mean duration to the preceding and subsequent biceps peaks were maximally close to the modal durations in each direction. To find sets of similar segments at movement onset, we first found the set of 50 segments that were 700 ms in length such that both biceps and triceps activity was below a minimally low threshold for the first 300 ms, and then their means over the last 400 ms were both above 0.1. From this set of 50, we then found the set of at least 30 segments such that the time from when biceps crosses the minimally low threshold to the first subsequent biceps peak and the time from when biceps crosses the minimally low threshold to the first subsequent triceps peak were both maximally close to their respective modal durations (among the set of 50). Here, modal durations were found as the peak of histograms with 5 ms bins. To find sets of similar segments at movement offset, we first computed a smoothed first derivative of the optical encoder signal reporting wheel position. An epoch of no wheel movement was identified in the resulting signal. A threshold was set as 5 times the standard deviation of the signal over this epoch. Time points where the signal rose above and fell below this threshold were identified. Segments where the signal was below threshold for less than ¼ of a second were redefined as being above threshold, as these did not reflect real stoppages in wheel turning, which is inherently iterative. We then identified all 700 ms segments where the signal was above threshold for 400 ms, then fell below threshold for 300 ms. We then ranked all segments both based on the mean of the signal over the first 400 ms and based on the mean of the summed biceps and triceps activity over the first 400 ms. We added these ranks together and found the 30 segments where the summed ranks were lowest.

Estimated spike time series for each dendritic cross-section (source) were aligned to muscle activity time series and Gaussian smoothed (10 ms standard deviation). Because the total number of sources was large and subsequent identification of rabies-labeled sources involved manual intervention, sources lacking substantial behaviorally correlated relevant activity were eliminated from further analysis if they did not have a substantial projection onto the top principal components of the trial-averaged activity of sources identified in the given image time series.

Sources from each image time series were classified as either rabies-labeled, unlabeled, or undetermined using a method that first identified the GCaMP6f image from the z stack collected following rabies injection with the highest correlation with the mean of each GCaMP6f image time series. The corresponding tdTomato image collected in the same plane was then used for subsequent source classification. Offsets between the selected GCaMP6f and tdTomato images were corrected by shifting one image relative to the other. The pixel masks for each source were then mapped to the selected GCaMP6f z stack image and outlines of pixel masks were defined for visual inspection and source classification. Sources were classified as rabies-labeled if (a) one cross-section in the GCaMP6f z stack image could be unambiguously linked to the source mask, and (b) that cross-section showed obvious tdTomato labeling above background. Sources were classified as unlabeled if (a) one cross-section in the GCaMP6f z stack image could be unambiguously linked to the source mask, but (b) that cross-section showed no discernable tdTomato labeling above background. Remaining sources were unclassified.

## SUPPLEMENTARY FIGURES

**Figure S1:**
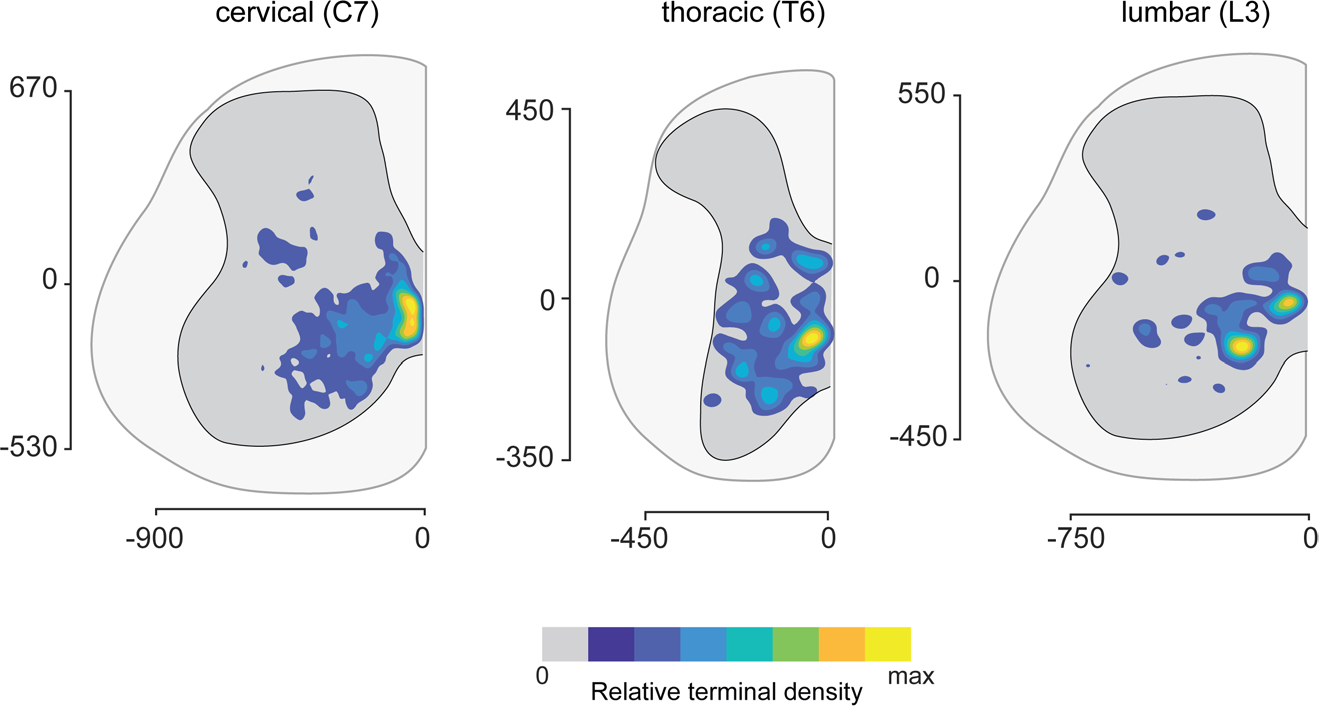
Ipsilateral corticospinal termination patterns at different rostrocaudal levels of the spinal cord Density maps of putative corticospinal terminations in the ipsilateral hemicord at three different spinal levels. These were generated using transverse spinal sections showing YFP-labeled corticospinal terminals after injection of *AAV2/1-Syn-hChR2-YFP* into ipsilateral motor and somatosensory cortex in adult mice, and co-localization with VGluT1. Axis labels are in microns.

**Figure S2:**
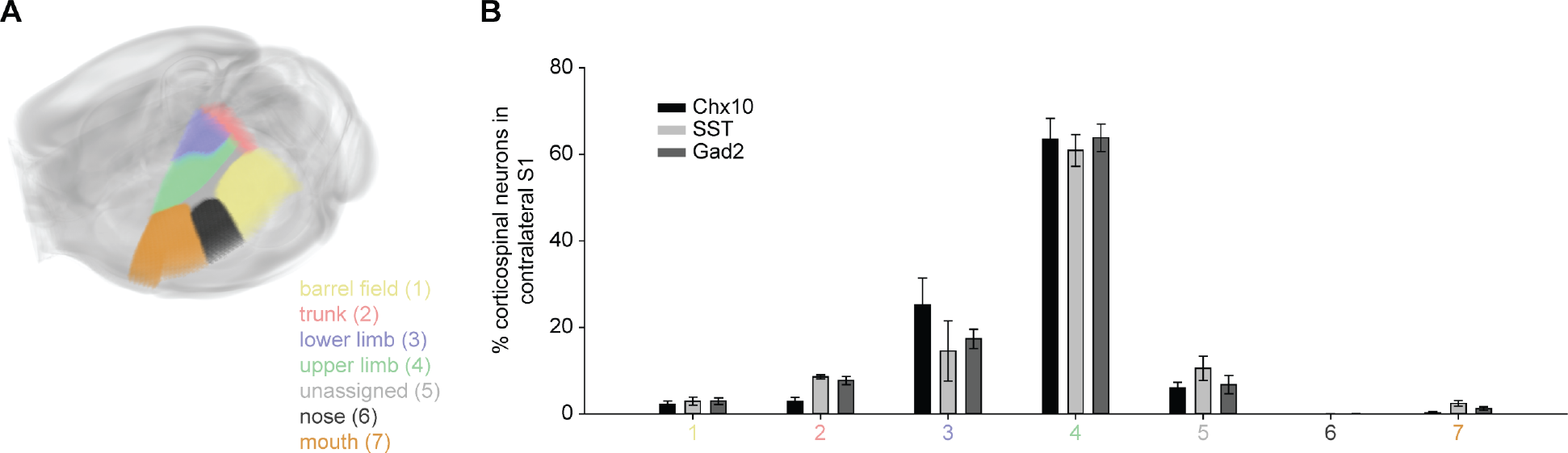
Forelimb S1 contains most corticospinal neurons following monosynaptic tracing from cervical spinal cord, demonstrating accuracy of registration (A) Schematic showing spatial extent of the topographic subdivisions of S1. (B) Mean ± s.e.m. percentage of labeled corticospinal neurons in S1 across its constituent subdivisions in each of the three genotypes examined.

**Figure S3:**
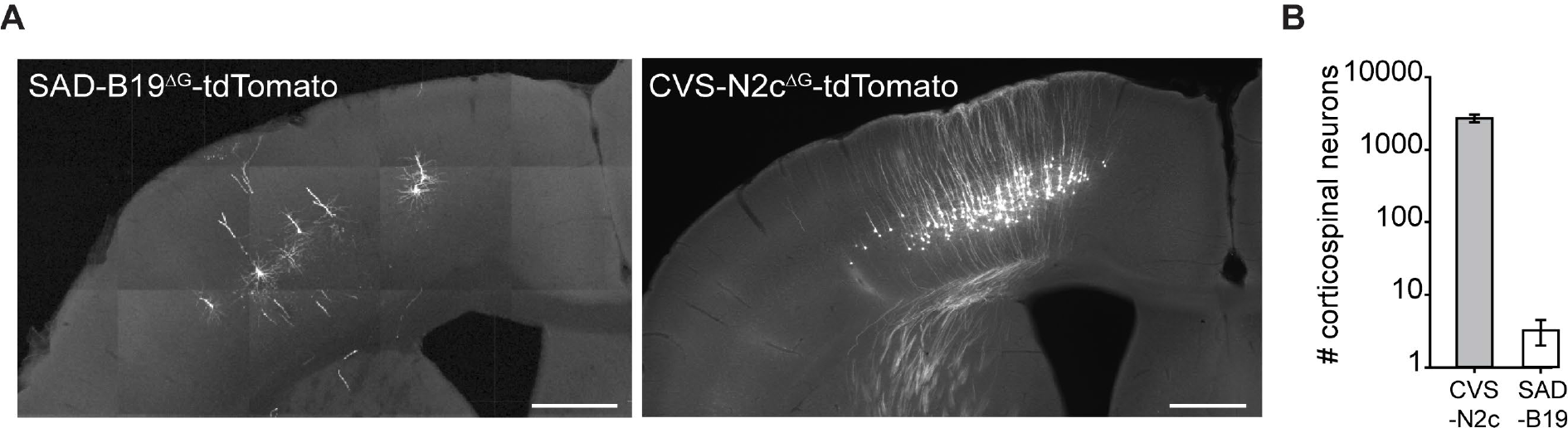
Enhanced efficiency of transsynaptic rabies tracing with CVS-N2c (A) Corticospinal neurons that formed monosynaptic contacts onto Chx10 expressing spinal interneurons were labeled following injection of tdTomato encoding pseudotyped rabies virus into cervical segments of *Chx10::Cre* mice. Corticospinal neurons were scarce with the SAD-B19^ΔG^ rabies variant (left panel) but efficiently labeled with the CVS-N2c^ΔG^ variant (right panel). Scale bars=500µm. (B) Mean ± s.e.m. number of labeled corticospinal neurons per animal across both hemispheres. The mean was ∼800 fold greater with CVS-N2c^ΔG^ rabies than with SAD-B19^ΔG^ rabies (n=4, p= 0.00023, two-tailed t-test). Data are combined from both *Chx10::Cre* and *Gad2::Cre* mice.

